# A NaV1.8^FlpO^ mouse enabling selective intersectional targeting of low threshold C fiber mechanoreceptors and nociceptors

**DOI:** 10.1101/2025.03.17.643556

**Authors:** John CY Chen, Lech Kaczmarczyk, Walker S Jackson, Max Larsson

## Abstract

Genetic targeting of select populations of cells in the mouse nervous system is often hampered by a lack of selectivity, as candidate genes for such targeting are commonly expressed by multiple cell populations, also in the same region. Intersectional targeting using two or more genes has been enabled by the development of reporter tools dependent on more than one recombinase or gene regulator. Still, widespread adoption of intersectional tools is complicated by a scarcity of driver mice expressing recombinases otherthan Cre. Here we report the generation and characterization of a new driver mouse that expresses the FlpO recombinase from the endogenous locus of the *Scn10a* gene encoding Na_V_1.8, a voltage-gated sodium channel that is almost exclusively expressed in the afferent limb of the peripheral nervous system. Moreover, among sensory neurons the channel is preferentially expressed in nociceptors and in low-threshold C-fiber mechanoreceptors (C-LTMRs). The mouse showed high recombination efficiency (97 %) and selectivity (93 %) in dorsal root ganglia. Reporter-expressing fibers were observed in a variety of peripheral tissues, including skin, skeletal muscle, genitalia, bladder and intestines. To validate the suitability of the FlpO mouse line for intersectional targeting, we crossed it with a mouse line expressing CreERT2 from the *Th* (tyrosine hydroxylase) locus. This approach resulted in strikingly selective and efficient targeting of C-LTMRs, showing robust visualization of nerve endings of these fibers in skin and spinal cord at the light and electron microscopic level. Thus, the Na_V_1.8^Flpo^ mouse line presented here constitutes a selective and versatile tool for intersectional genetic targeting of Na_V_1.8 expressing primary afferent neurons.

## Introduction

Primary afferent fibers innervating skin and internal organs are pivotal for sensing the external and internal environments, thus informing behaviour and homeostasis that enable the survival of the organism. Such sensory fibers include low-threshold mechanoreceptors (LTMRs) that signal innocuous mechanical stimuli, as well as nociceptors that are activated by noxious stimuli that threaten the integrity of the tissue. The last decade has seen a rapid progress in delineating primary afferent populations via transcriptomics and other means, identifying genetic markers of such populations (e.g., Kupari and Ernfors 2023; Sharma et al. 2020; Usoskin et al. 2015; Handler and Ginty 2021; Li et al. 2015). However, whereas in some cases this has allowed targeting of relatively homogeneous populations for functional and anatomical characterization using Cre-expressing mouse lines (Qi et al. 2024b; Li et al. 2011), such an approach is often hampered by the lack of a single genetic marker that uniquely identifies the population of interest.

C fiber low-threshold mechanoreceptors (C-LTMRs) are thought to mediate pleasant touch (Löken et al. 2009; Olausson et al. 2002; Huzard et al. 2022) but have also been suggested to have a role in pain modulation (Larsson and Nagi 2022). However, investigation of the function of these fibers in animals has been hindered by a lack of genetic tools for selective functional manipulation. C-LTMRs express *Th* (encoding tyrosine hydroxylase) and *Slc17a8* (encoding the vesicular glutamate transporter 3, VGluT3); however, those genes are also expressed in other cells in the CNS and peripheral tissue, which complicates the use of mice expressing Cre or tamoxifen-inducible CreERT2 from the loci of either of these genes. For instance, targeting of a genetically encoded actuator protein using *Th*^CreERT2^ mice may also capture mechanosensitive Merkel cells in the skin (Hoffman et al. 2018), sympathetic nerve fibers, as well as catecholaminergic neurons in the brainstem. However, unlike those cells, C-LTMRs also express the voltage-sensitive sodium channel Na_V_1.8, encoded by the gene *Scn10a*. Thus, it may be possible to employ an intersectional targeting approach that uses *Th*^CreERT2^ driver mice together with a second driver mouse line expressing a distinct recombinase from the *Scn10a* locus, in combination with double recombinase dependent genetic tools. Moreover, in addition to C-LTMRs, Na_V_1.8 is selectively expressed in most nociceptors whereas myelinated LTMRs have little or no expression of this ion channel. A Na_V_1.8 driver mouse could therefore potentially be used for intersectional targeting of nociceptor subpopulations where the second genetic marker is not nociceptor-specific. We therefore generated a new mouse line that expresses FlpO recombinase from the *Scn10a* locus and explored its utility for targeting of Na_V_1.8-expressing primary afferent neurons in somatosensory and viscerosensory pathways, including intersectional selective targeting of C-LTMRs.

## Material and methods

### Gene targeting

The Na_V_1.8^FlpO^ knock-in mouse line was generated through homologous recombination in J1 embryonic stem cells (derived from 129S4 mice) at the Karolinska Center Transgene Technology. The Scn10a-IRES2-FlpO targeting vector (TV) was constructed using the Gibson Assembly cloning kit (New England Biolabs). This vector incorporated a self-excision tACE-Cre-Neo (ACN) selection cassette, similar to the design originally described by Bunting et al. (1999), but with Lox2272 flanking sites. The plasmid containing the ACN cassette was kindly provided by Mario Capecchi (Addgene plasmid #20343; Wu et al. 2008). Homology arms for the targeting vector were amplified via PCR from mouse ES cell DNA using the following primers: Forward: CCCGTGTGCCAGGAACTGAGTC Reverse: CATCAACTCATGCTTGGATGGTGT. The vector backbone was based on the Slc1a3-CreERT2 plasmid (Addgene plasmid #129409; Kaczmarczyk et al. 2021), linearized with AscI and SmaI restriction enzymes. To enhance homologous recombination efficiency, the TV was co-electroporated with a pX330-Nav1.8 Cas9 vector containing a suitable sgRNA sequence (ATGCTGGAGTGTCTTCACTG). The pX330 backbone was a gift from Feng Zhang (Addgene plasmid #42230; Cong et al. 2013). The full sequences of both the TV and the Cas9 plasmid are available at https://tinyurl.com/Scn10a-IRES2-FlpO. Plasmid integrity was verified by restriction digest and Sanger sequencing. As a result of successful gene targeting, the Scn10a locus encoding Na_V_1.8 was replaced with an Scn10a-IRES2-FlpO-3’UTR cassette (Fig. 1).

**Figure 1.**
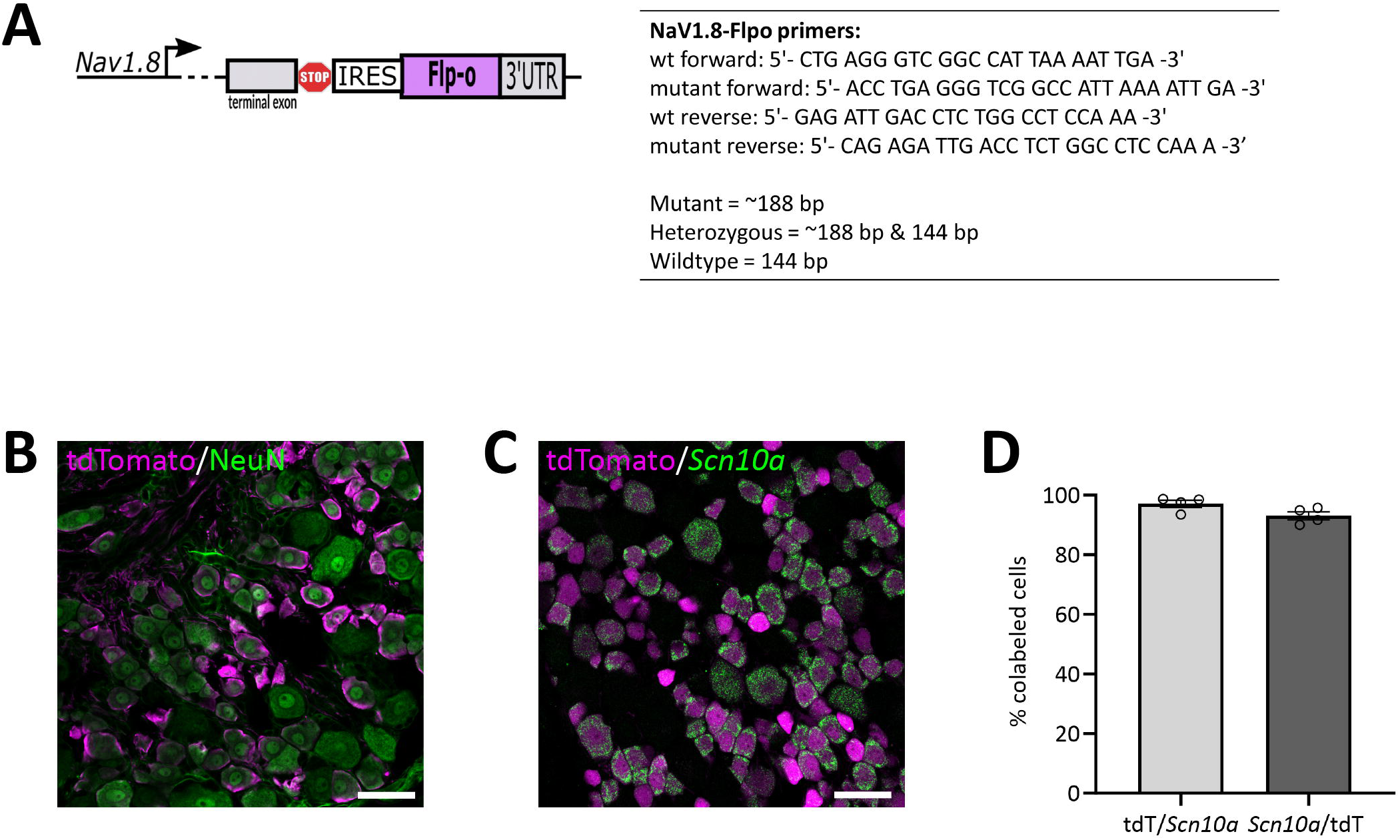
Verification of the Na_V_1.8^FlpO^ mouse line. **A**, Design of the allele and genotyping primers. **B**, expression of tdTomato in NeuN^+^ neurons in a lumbar DRG of a Na_V_1.8^Flpo^;Ai65F mouse. **C**, co-localization of tdTomato with *Scn10a* transcript in a lumbar DRG. **D**, Recombination efficiency with respect to *Scn10a*^+^ cells (mean ± S.E.M.) in L3-L5 DRGs. Each data point indicates a section from a single ganglion; n=3 mice. Scale bars, 50 µm.

### Animals

For Flp-dependent reporter expression, Na_V_1.8^FlpO^ mice were crossed with Ai65F mice (RCF-tdT; Jackson Laboratory, strain #032864) (Daigle et al. 2018). For intersectional genetic targeting of C-LTMRs, Na_V_1.8^FlpO^ mice were crossed with either Ai65 mice (RCFL-tdT; Jackson Laboratory, strain #021875) (Daigle et al. 2018) or ROSA26DR-Matrix-dAPEX2 (Jackson Laboratory, strain #032764; here called APEX2 mice) (Zhang et al. 2019); the resulting offspring was further crossed with *Th*^CreERT2^ mice (Jackson Laboratory, strain #025614) (Abraira et al. 2017) to generate offspring heterozygous for Na_V_1.8^FlpO^, *Th*^CreERT2^, and reporter. *Th*^CreERT2^ mice were also crossed with singly Cre-dependent Ai14 mice (RCL-tdTomato, Jackson Laboratory, strain #007914) (Madisen et al. 2010). Mouse genotyping was performed using the following primers: wt forward, 5’-CTG AGG GTC GGC CAT TAA AAT TGA-3’; mutant forward, 5’-ACC TGA GGG TCG GCC ATT AAA ATT GA-3’; wt reverse, 5’-GAG ATT GAC CTC YGG CCT CCA AA-3’; mutant reverse, 5’CAG AGA TTG ACC TCT GGC CTC CAA A-3’. All animal experiments were approved by the Animal Ethics Committee at Linköping University (permit no. 2439-2021) and performed in accordance with the EU Directive 2010/63/EU.

### Tamoxifen injection

For *Th*^CreERT2^;Na_V_1.8^FlpO^;Ai65 and *Th*^CreERT2^;Na_V_1.8^FlpO^;APEX2 mice, tamoxifen (T5648, Sigma-Aldrich) dissolved in corn oil at a concentration of 20 mg/mL was administered at 8-10 weeks of age by a single injection (100 µL, i.p.). The mice were perfused no earlier than two weeks after injection.

### Tissue preparation

Adult mice of either sex were anesthetized with i.p. injections of either sodium pentobarbital (100 mg/kg) or a mixture of ketamine (120 mg/kg) and dexmedetomidine (0.5 mg/kg), after which they were subjected to transcardial perfusion using 5 mL phosphate buffer (PB, 0.1 M pH 7.4) followed by 50 mL of 4 % paraformaldehyde (for immunofluorescence) or 2 % paraformaldehyde and 2.5 % glutaraldehyde (for electron microscopy). Tissue of interest was harvested, post-fixed overnight at 4°C and then stored in 1/10 fixative or PB until further processing.

### Immunofluorescence

Tissue specimens were cryoprotected in 30 % sucrose, embedded in OCT, cut at 15 µm or 50 µm thickness in a cryostat and placed on glass slides. Alternatively, sections were embedded in 4 % low-melting agarose and cut on a vibrating microtome (Campden Instruments 7000smz-2) at 50 µm thickness. Sections were incubated in phosphate-buffered saline (PBS) with 3 % normal goat serum, 0.5 % bovine serum albumin and 0.5 % Triton X-100 (blocking solution), and then in blocking solution with primary antibodies (see Table 1) overnight at room temperature. Sections were next rinsed and incubated in secondary antibody solution (Table 2). For detection of isolectin B_4_ (IB_4_) binding sites, biotinylated IB_4_ (Life Technologies, I21214) was added to the primary antibody solution at 1:1000 dilution, and streptavidin-Alexa Fluor 488 (Life Technologies, S11223) added to the secondary antibody solution at 1:500 dilution. In some cases, cell nuclei were stained using DAPI or SYTOX Deep Red (ThermoFisher Scientific, S11380). Sections were coverslipped with Prolong Diamond, Prolong Glass or SlowFade Diamond (Life Technologies) and imaged using a Zeiss LSM800 or Leica Stellaris 5 confocal microscope.

**Table 1.** Primary antibodies.

| **Antigen** | **Host, isotype** | **Supplier** | **Cat #** | **Dilution** |
| --- | --- | --- | --- | --- |
| CGRP | Guinea pig | Synaptic Systems | 414 004 | 1:1000 |
| NFH | Chicken | Thermo Fisher Scientific | PA1-10002 | 1:1000 |
| NeuN | Mouse, IgG<sub>1</sub> | Merck Millipore | MAB377 | 1:1000 |
| Parvalbumin | Guinea pig | Synaptic Systems | 195 004 | 1:500 |
| PKCγ | Guinea pig | Frontier Institute | PKCγ-GP-Af350 | 1:250 |
| RFP (tdTomato) | Chicken | Synaptic Systems | 409 006 | 1:2000 |
| RFP (tdTomato) | Rabbit | Rockland | 600-401-379 | 1:1000 |
| TH | Rabbit | Abcam | Ab112 | 1:200 |
| TRPV1 | Rabbit | Synaptic Systems | 444 033 | 1:1000 |
| VGLUT3 | Guinea pig | Frontier Institute | VGLUT3-GP-Af300 | 1:500 |

**Table 2.** Secondary antibodies.

| **Host, target** | **Fluorophore** | **Supplier** | **Cat #** | **Dilution** |
| --- | --- | --- | --- | --- |
| Goat α-chicken IgY (H+L) | Alexa Fluor 488 | Life Technologies | A32931 | 1:500 |
| Goat α-chicken IgY (H+L) | Alexa Fluor 555 | Life Technologies | A21437 | 1:500 |
| Goat α-guinea pig IgG (H+L) | Alexa Fluor 488 | Life Technologies | A11073 | 1:500 |
| Goat α-guinea pig IgG (H+L) | Alexa Fluor 647 | Life Technologies | A21450 | 1:500 |
| Goat α-mouse IgG<sub>1</sub> | Alexa Fluor 488 | Life Technologies | A21121 | 1:500 |
| Goat α-rabbit IgG (H+L) | Alexa Fluor 488 | Life Technologies | A11034 | 1:500 |
| Goat α-rabbit IgG (H+L) | Alexa Fluor Plus 555 | Life Technologies | A32732 | 1:500 |

### Electron microscopy

The day after perfusion fixation, thoracic spinal cord from three *Th*^CreERT2^;Na_V_1.8^FlpO^;APEX2 mice was embedded in 4 % low-melting agarose, cut on a vibrating microtome into 50 µm or 150 µm transverse sections, and immediately subjected to APEX2 histochemistry. The sections were incubated in 3,3’-diaminobenzidine (DAB) without H_2_O_2_ for 30-60 min, and then in DAB with H_2_O_2_ (Vector Laboratories SK-4100) for 45-60 min. The sections were then osmicated in 1 % OsO_4_ in 0.1 M PB, stained with 1 % uranyl acetate in 70 % ethanol, dehydrated in a graded series of ethanol and embedded in Durcupan. Ultrathin sections were placed on Formvar coated single-slot copper grids, counterstained with 2 % aqueous uranyl acetate and 0.5 % lead citrate (in some cases post-embedding counterstaining was omitted) and examined in a JEOL JEM1400 Flash transmission electron microscope at 80 kV accelerating voltage.

### Statistical analysis

All quantitative data were prepared in GraphPad Prism and are given as mean ± S.E.M.

## Results

Mice heterozygous or homozyogous for the *Scn10a*-IRES2-*FlpO* allele were viable, fertile and showed no overt behavioral deficits. In lumbar DRGs of Na_V_1.8^FlpO^;Ai65F mice, tdTomato protein was found in numerous NeuN immunoreactive neurons of different sizes (Fig. 1). *In situ* hybridization showed that the vast majority (97.1 ± 1.2 %) of *Scn10a*^+^ neurons expressed tdTomato protein; conversely, 93.1 ± 1.4 % of tdTomato^+^ cells expressed *Scn10a* mRNA. Calcitonin gene-related peptide (CGRP), IB_4_ binding, and TRPV1 were found in subsets of tdTomato^+^ neurons (Fig. 2). tdTomato also showed partial overlap with neurofilament heavy chain (NFH) and its transcript *Nefh*, including in peripheral axons of the sciatic nerve. In the spinal cord, tdTomato^+^ processes, i.e. presumed central projections of Na_V_1.8^+^ primary afferent fibers, were present throughout the dorsal horn at varying densities (Fig. 3). The largest density of tdTomato^+^ processes was found in lamina I and the dorsal half of lamina II, whereas ventral lamina II and dorsal lamina III, as outlined by the presence of PKCγ^+^ neurons, showed a progressively lower density of processes along the dorsoventral axis. Processes were at all spinal levels considerably sparser in ventral lamina III to lamina V, and absent from the intermediate or ventral horns. Cell bodies exhibiting tdTomato fluorescence were not observed in the spinal cord. In spinal white matter, processes were found in Lissauer’s tract as well as in the dorsal and medial parts of the dorsal funiculus. We did not attempt a comprehensive mapping of the distribution of tdTomato^+^ cells and processes in the brain. However, tdTomato^+^ structures were generally sparse (Fig. 4). In the brainstem, tdTomato^+^ processes were found in the trigeminal spinal nucleus, as well as in the nucleus of the solitary tract. Notably, fibers were also present in the gracile nucleus (and to a lesser degree the cuneate nucleus), as well as in the principal trigeminal nucleus, the superior olivary complex, and in pontine nuclei. In the subcortical forebrain, tdTomato^+^ cells were noted in the anterior olfactory nucleus, the bed nucleus of the stria terminalis, the endopiriform nucleus, the septum and the nucleus of the diagonal band. In the cortex, tdTomato^+^ cells were generally not found; however, in frontal cortex we observed some tdTomato^+^ cells in layer 6b immediately superior to the corpus callosum. In addition, tdTomato^+^ fibers were quite abundant in layer 1a of the frontal cortex, whereas such fibers were very sparse in other cortical regions. Scattered tdTomato^+^ cells were found in the granule cell layer of the olfactory bulb.

**Figure 2.**
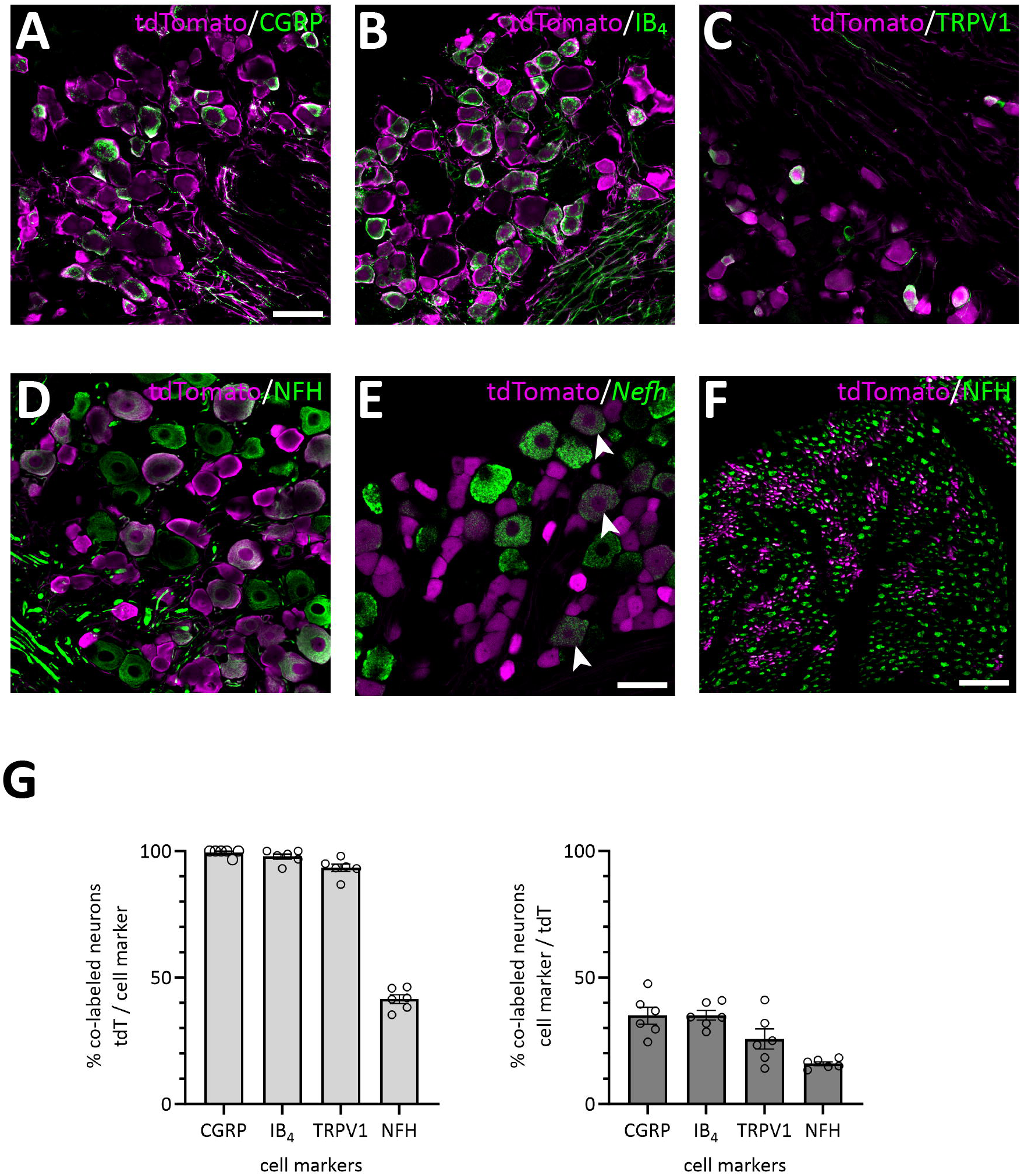
Primary afferent populations targeted by the Na_V_1.8^FlpO^ mouse line. **A-E**, Co-localization of tdTomato immunoreactivity (IR) with CGRP-IR, IB4 binding, TRPV1-IR, NFH-IR and *Nefh* transcript in lumbar DRGs. Scale bar in A, 50 µm, valid for A-D; scale bar in E, 50 µm. **F**, Co-localization of tdTomato-IR and NFH-IR in sciatic nerve. Scale bar, 25 µm. **G**, quantification of the co-localization of tdTomato with the indicated cell markers. Error bars indicate S.E.M.

**Figure 3.**
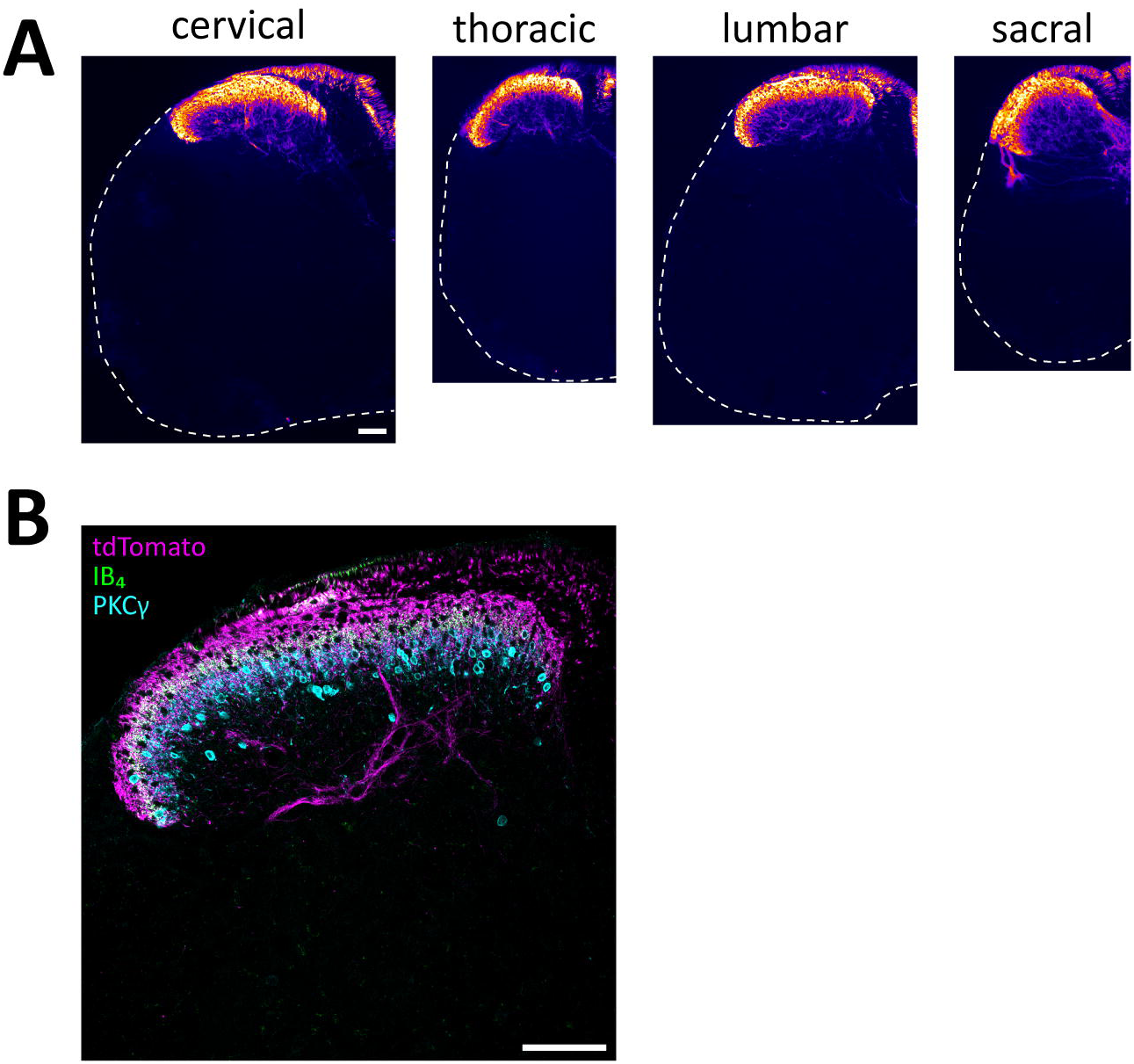
tdTomato^+^ processes in the spinal cord of Na_V_1.8^FlpO^;Ai65F mice. **A**, distribution of tdTomato^+^ processes in the spinal cord at different spinal levels. The fluorescence is false-colored to better visualize sparse processes. Dashed lines outline section edges where fluorescence is absent. Scale bar is 100 µm, valid for all panels. **B**, laminar distribution of tdTomato^+^ fibers in relation to IB_4_ binding and PKCγ immunolabeling. Scale bar, 100 µm.

**Figure 4.**
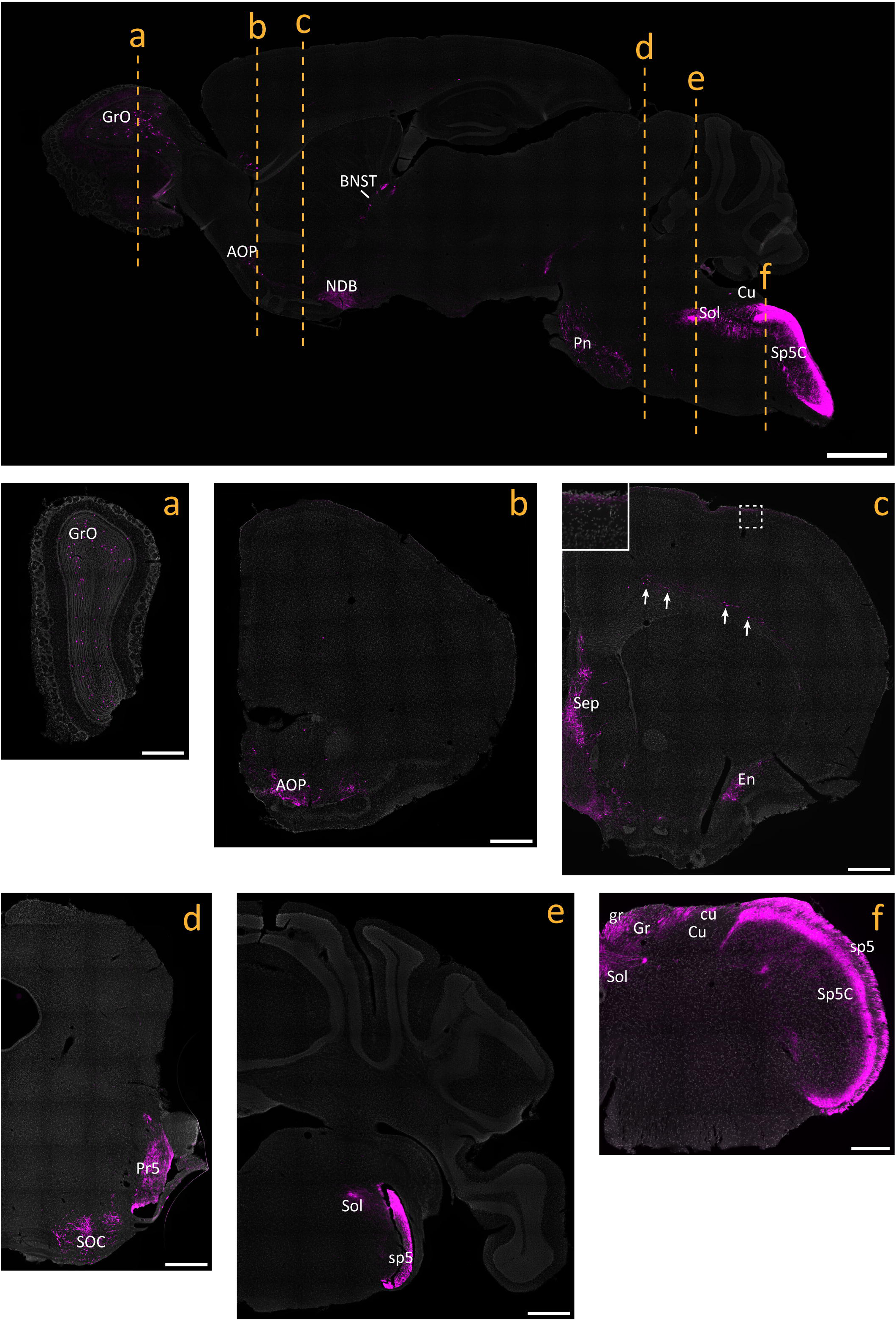
tdTomato^+^ processes and cells in the brain of Na_V_1.8^FlpO^;Ai65F mice. Top panel shows a parasagittal brain section. Center and bottom panels show coronal sections obtained at the levels indicated in the parasagittal section. All panels are counterstained with SYTOX Deep Red. Inset in c shows the portion indicated by dashed frame. Arrows indicate tdTomato^+^ neurons in layer 6 of frontal cortex. AOP, anterior olfactory nucleus, posterior part; BNST, bed nucleus of the stria terminalis; cu, cuneate fascicle; Cu, cuneate nucleus; En, endopiriform nucleus; gr, gracile fascicle; Gr, gracile nucleus; GrO, granular layer of the olfactory bulb; NDB, nucleus of the diagonal band; Pn, pontine nuclei; Pr5, principal sensory trigeminal nucleus; Sep, septal region; SOC, superior olivary complex; Sol, nucleus of the solitary tract; sp5, trigeminal tract; Sp5C, caudal spinal trigeminal nucleus. Scale bar in parasagittal section, 1 mm; scale bars in a-e, 500 µm; scale bar in f, 250 µm.

Select peripheral organs and ganglia were surveyed for the presence of tdTomato^+^ nerve fibers. In glabrous skin, tdTomato^+^ fibers forming free epidermal nerve endings were abundant (Fig. 5A). Notably, although apically directed fibers traversed dermal papillae, we observed no Meissner corpuscle-like endings, as has previously been reported in Na_V_1.8^Cre^ mice (Shields et al. 2012). In hairy skin, tdTomato^+^ circumferential nerve endings around hair follicles were also observed in addition to free nerve endings (Fig. 5B). In both glabrous and hairy skin, cutaneous tdTomato^+^ fibers often but not always colocalized with CGRP immunoreactivity.

**Figure 5.**
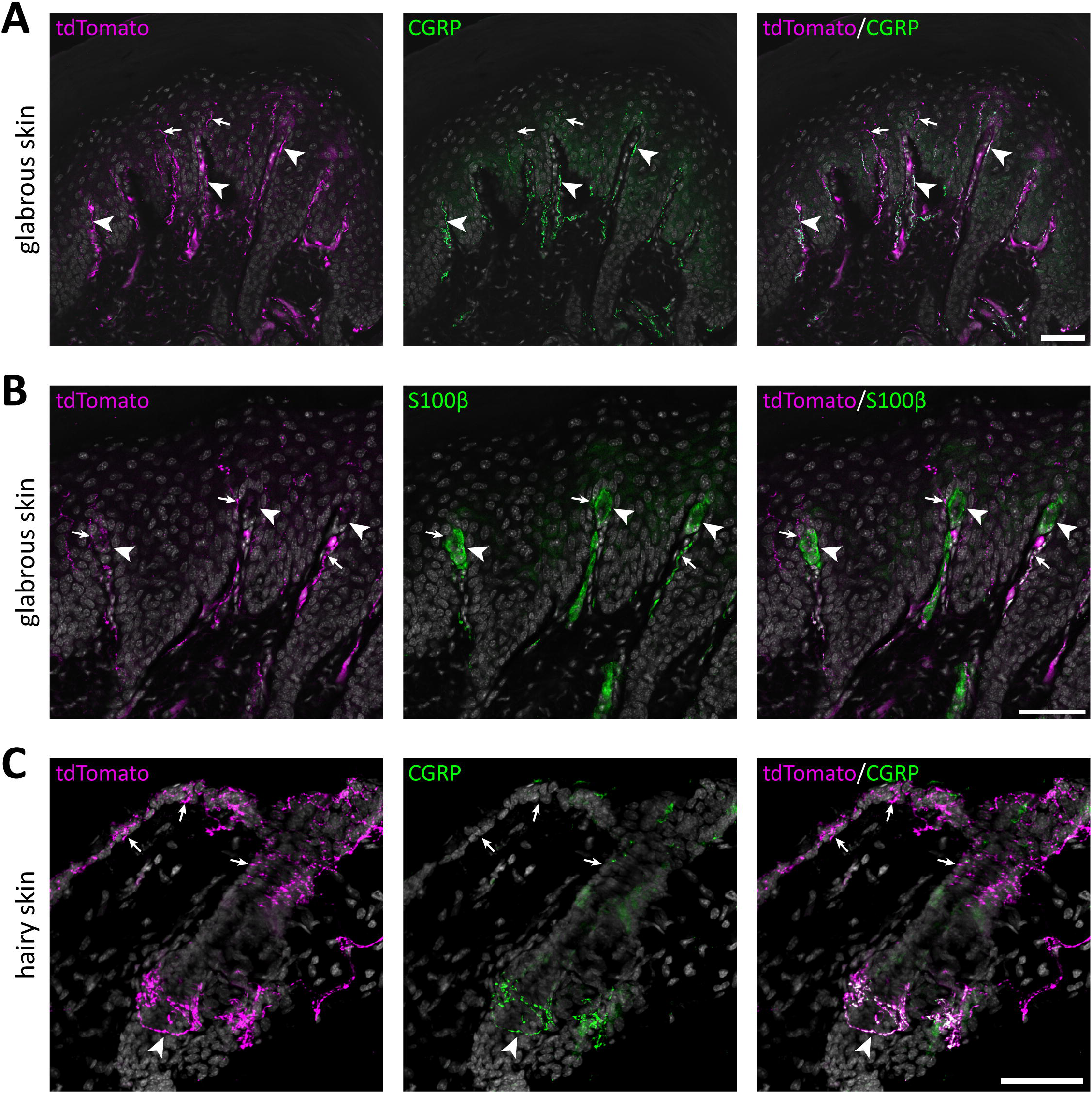
tdTomato^+^ processes in the skin of of Na_V_1.8^FlpO^;Ai65F mice. **A**, tdTomato^+^ processes and co-localization with CGRP-IR in hind paw glabrous skin. tdTomato^+^ fibers were frequently CGRP-IR in the dermis, whereas tdTomato^+^ fibers that lacked CGRP-IR extended into the epidermis. Arrowheads indicate examples of tdTomato^+^/CGRP^+^ fibers, and arrows indicate tdTomato^+^/CGRP^-^ fibers. Scale bar, 50 µm. **B**, Co-immunolabeling of tdTomato and S100β in hind paw glabrous skin. Whereas tdTomato^+^ fibers often travelled in dermal papillae, they never formed bundles associated with S100β^+^ Meissner corpuscles. Arrowheads indicate Meissner corpuscles; arrows indicate examples of tdTomato^+^ fibers juxtaposed to Meissner corpuscles. Scale bar, 50 µm. **C**, tdTomato^+^ fibers and co-localization with CGRP in hairy back skin. tdTomato^+^ fibers formed circumferential CGRP-IR nerve endings around hair follicles (arrowhead), but also free nerve endings in the epidermis that generally lacked CGRP immunoreactivity (arrows). Scale bar, 50 µm. All micrographs are counterstained with DAPI.

Many cells in nodose and jugular ganglia exhibited tdTomato immunoreactivity, as did neurons in the Grüneberg ganglion, a chemo-/thermosensory ganglion in the nose (Fig. 6). In skeletal muscle, sparse tdTomato^+^ fibers were found, while epithelia of the bladder, colon and rectum were richly innervated by tdTomato^+^ fibers. Similarly, tdTomato^+^ fibers were abundant in epithelia of both male and female genitalia. In the aortic arch, sparse tdTomato^+^ fibers were detected, whereas the lamina propria of the olfactory mucosa showed strong innervation by tdTomato^+^ fibers.

**Figure 6.**
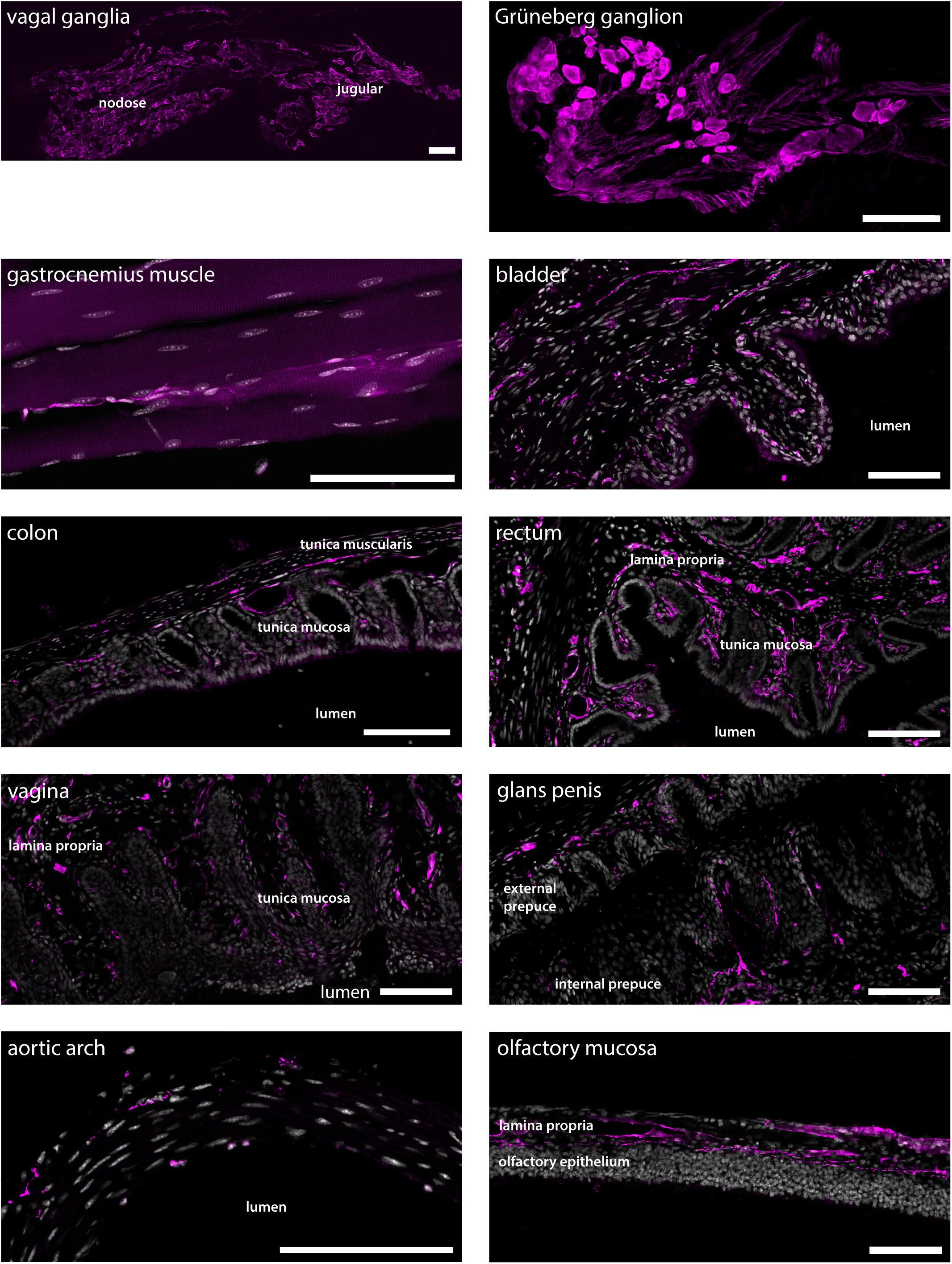
tdTomato^+^ processes and cells in non-dermal peripheral tissue. Scale bars are 100 µm in all panels.

In order to test the utility of the Na_V_1.8^FlpO^ strain for intersectional genetic targeting of primary afferent fiber populations, we crossed this line with a *Th*^CreERT2^ mouse strain and a Cre/Flp-dependent reporter line (Ai65). In DRGs of these mice, 75.3 ± 1.8 % of *Th*^+^ neurons showed tdTomato fluorescence, whereas 94.3 ± 1.7 % of tdTomato^+^ cells were *Th*^+^, indicating high recombination efficiency and selectivity towards C-LTMRs among primary afferent fibers (Fig. 7). Importantly, tdTomato fluorescence did not co-localize with nociceptive markers CGRP or IB4, or with the myelinated neuron marker NFH (Fig. S1). In the lumbar enlargement of the spinal cord of these mice, tdTomato^+^ processes were evident in a distinct band in the superficial dorsal horn. This band coincided with the dorsal part of the region occupied by neurons expressing the γ isoform of protein kinase C (PKCγ), ventral to the plexus formed by IB_4_ binding C-fibers (Fig. 8A). Notably, tdTomato^+^ fibers were absent from the medial dorsal horn that receives input from hind paw glabrous skin, and co-localized with VGLUT3^+^ structures. In *Th*^CreERT2^;Na_V_1.8^FlpO^;Ai65 mice, tdTomato fluorescence was restricted to the dorsal roots, tract of Lissauer and inner lamina II, whereas in *Th*^CreERT2^;Ai14 mice that target tdTomato to all TH^+^ cells, various tdTomato^+^ processes were also observed throughout gray and white matter (Fig. 8B). In the hairy skin of *Th*^CreERT2^;Na_V_1.8^FlpO^;Ai65 mice, tdTomato fluorescence was found selectively in lanceolate endings surrounding hair follicles, whereas in Th^CreERT2^;Ai14 mice tdTomato fluorescence was also observed in presumed sympathetic fibers. In the brain of *Th*^CreERT2^;Na_V_1.8^FlpO^;Ai65 mice tdTomato^+^ processes were restricted to the solitary tract, the nucleus of the solitary tract, and, very sparsely, trigeminal structures; other parts of the brain were completely devoid of tdTomato^+^ processes. By contrast, tdTomato^+^ fibers and cells showed the expected widespread distribution in *Th*^CreERT2^;Ai14 mice (Fig. S2). Thus, using the Na_V_1.8FlpO strain affords highly efficicent and selective targeting of C-LTMRs while avoiding targeting of central and peripheral catecholaminergic systems.

**Figure 7.**
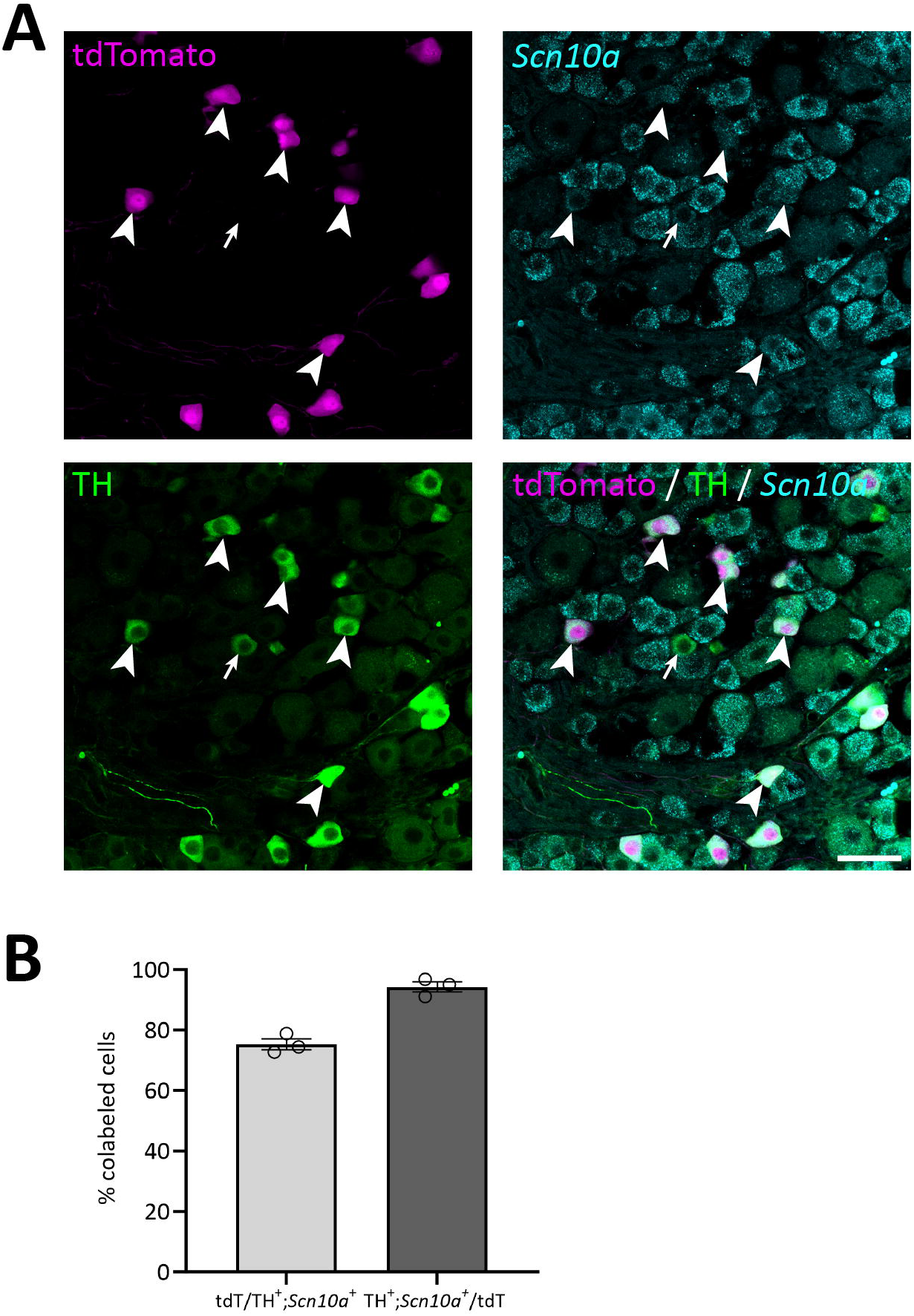
Verification of the *Th*^CreERT2^;Na_V_1.8^FlpO^;Ai65 mouse line. **A**, Section of a lumbar DRG subjected to immunolabeling of tdTomato and TH, and to *in situ* hybridization for *Scn10a*. Arrowheads indicate cells triple positive for tdTomato, TH and *Scn10a*. Arrow indicates an example of a TH^+^/*Scn10a*^+^ cell that lacks tdTomato expression. Scale bar, 50 µm. **B**, Quantification of recombination efficiency with respect to TH^+^/*Scn10a*^+^ cells. Note that all TH^+^ cells were *Scn10a*^+^ and vice versa. Each data point corresponds to a single DRG section and mouse; n = 3 mice. Error bars indicate S.E.M.

**Figure 8.**
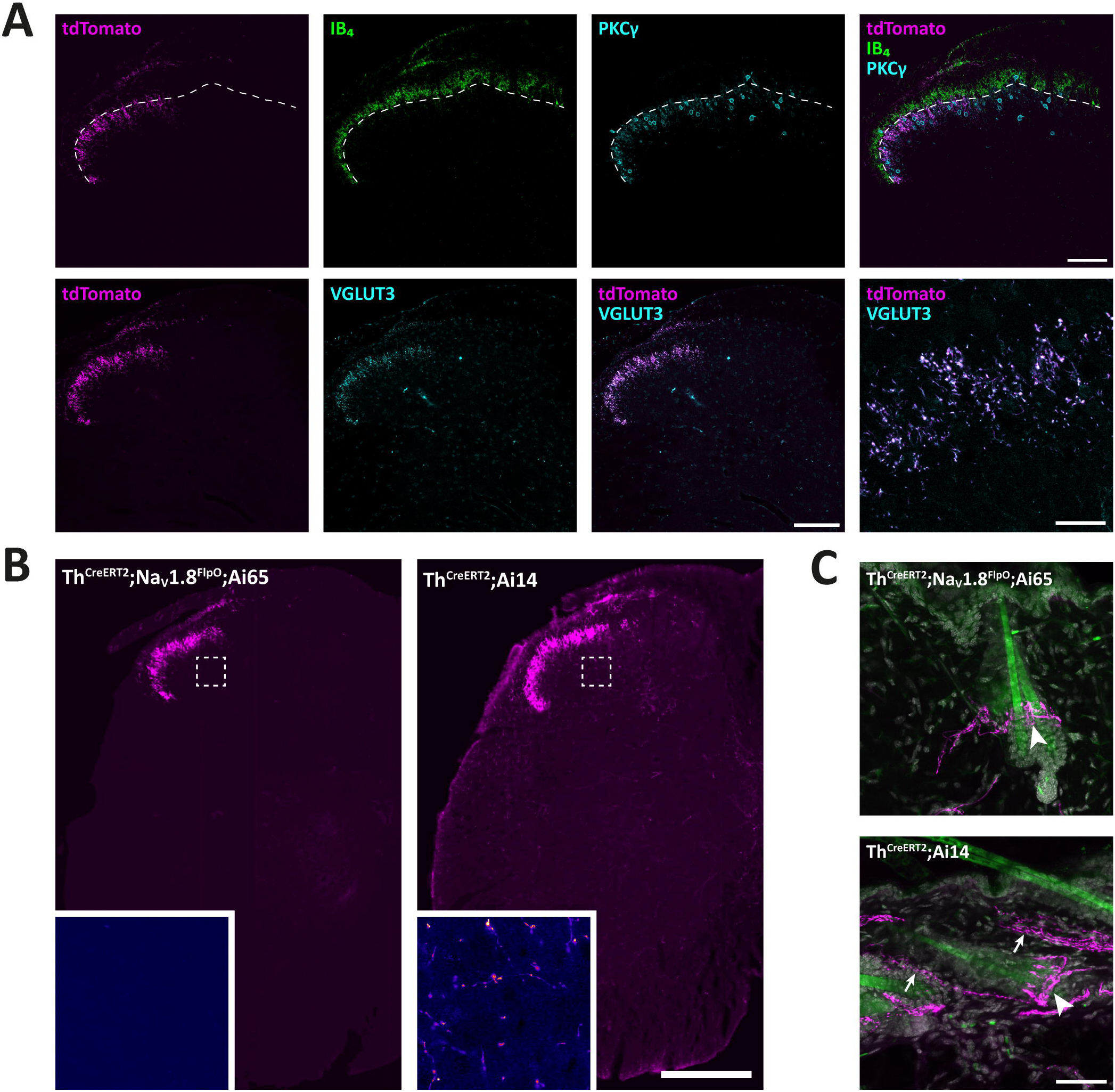
Targeting of C-LTMRs in spinal cord and skin of *Th*^CreERT2^;Na_V_1.8^FlpO^;Ai65 mice. **A**, distribution of tdTomato^+^ processes in the superficial dorsal horn of L4 spinal cord with respect to IB_4_ binding and PKCγ and VGluT3 immunolabeling. Dashed line indicates ventral border of the IB_4_ plexus. Note that the band of tdTomato^+^ fibers and endings were almost entirely ventral to this border, and localized to the dorsal portion of the region inhabited by PKCγ neurons. Note that the medial gray matter receiving input from glabrous skin was devoid of tdTomato^+^ fibers. tdTomato^+^ processes showed near-complete co-localization with VGluT3 in lamina II. Scale bars are 100 µm except in bottom-rightmost panel, which is 20 µm. **B**, Comparison of spinal distribution of tdTomato^+^ processes in *Th*^CreERT2^;Na_V_1.8^FlpO^;Ai65 versus *Th*^CreERT2^;Ai14 mice. In the former, tdTomato^+^ processes were only observed in the superficial dorsal horn as well as in the dorsal root and Lissauer’s tract, whereas in *Th*^CreERT2^;Ai14 mice tdTomato^+^ processes were present throughout gray and white matter. Insets show magnified false-color views of the regions in the deep dorsal horn indicated by dashed frames. Scale bar, 250 µm. **C**, Hairy skin innervation by tdTomato^+^ fibers in *Th*^CreERT2^;Na_V_1.8^FlpO^;Ai65 versus *Th*^CreERT2^;Ai14 mice. tdTomato^+^ lanceolate nerve endings presumably formed by C-LTMRs were found around hair follicles in both mice (indicated by arrowheads). In *Th*^CreERT2^;Ai14 mice presumed sympathetic nerve fibers were also tdTomato^+^ (arrows). Scale bar, 50 µm.

The ultrastructure of central C-LTMR terminations is contentious, having been described as forming either type I or type II glomeruli in the spinal dorsal horn (Larsson and Broman 2019; Salio et al. 2021). Such discrepancy could be partly attributed to differences in the markers used for C-LTMR terminals. Here we addressed this issue by creating *Th*^CreERT2^;Na_V_1.8^FlpO^;APEX2 mice, expressing APEX2 in the mitochondrial matrix of C-LTMRs in a Cre-/Flp-dependent manner highly selective for C-LTMRs. In spinal cord from these mice, terminals possessing mitochondria with APEX2 reaction product were commonly found to constitute central terminals of synaptic glomeruli in inner lamina II (Fig. 9). Often irregularly shaped, these terminals exhibited light axoplasm and loosely packed clusters of round and clear synaptic vesicles. In many cases, multiple asymmetric synapses were formed with postsynaptic dendrites. Some of these dendrites contained vesicles and thus presumably originated from inhibitory interneurons (Todd 1996). Very occasionally, a presynaptic axon was found to form a symmetric synapse onto the central APEX2^+^ terminal (Fig. 9B). Sometimes, the central terminal exhibited more tightly packed vesicle clusters and a scalloped shape, somewhat reminiscent of the central terminals of type I glomeruli (Fig. 9C). However, the central terminals of type I glomeruli exhibited a distinctly more electron dense axoplasm and were considerably more tightly packed with vesicles which were of irregular sizes, unlike the more homogenously sized vesicles of APEX2^+^ terminals. Notably, type I glomeruli were almost always more dorsally located than APEX2^+^ terminals. Thus, C-LTMRs form type II glomeruli that are readily distinguishable from type I glomeruli formed by IB_4_ binding non-peptidergic C fiber nociceptors.

**Figure 9.**
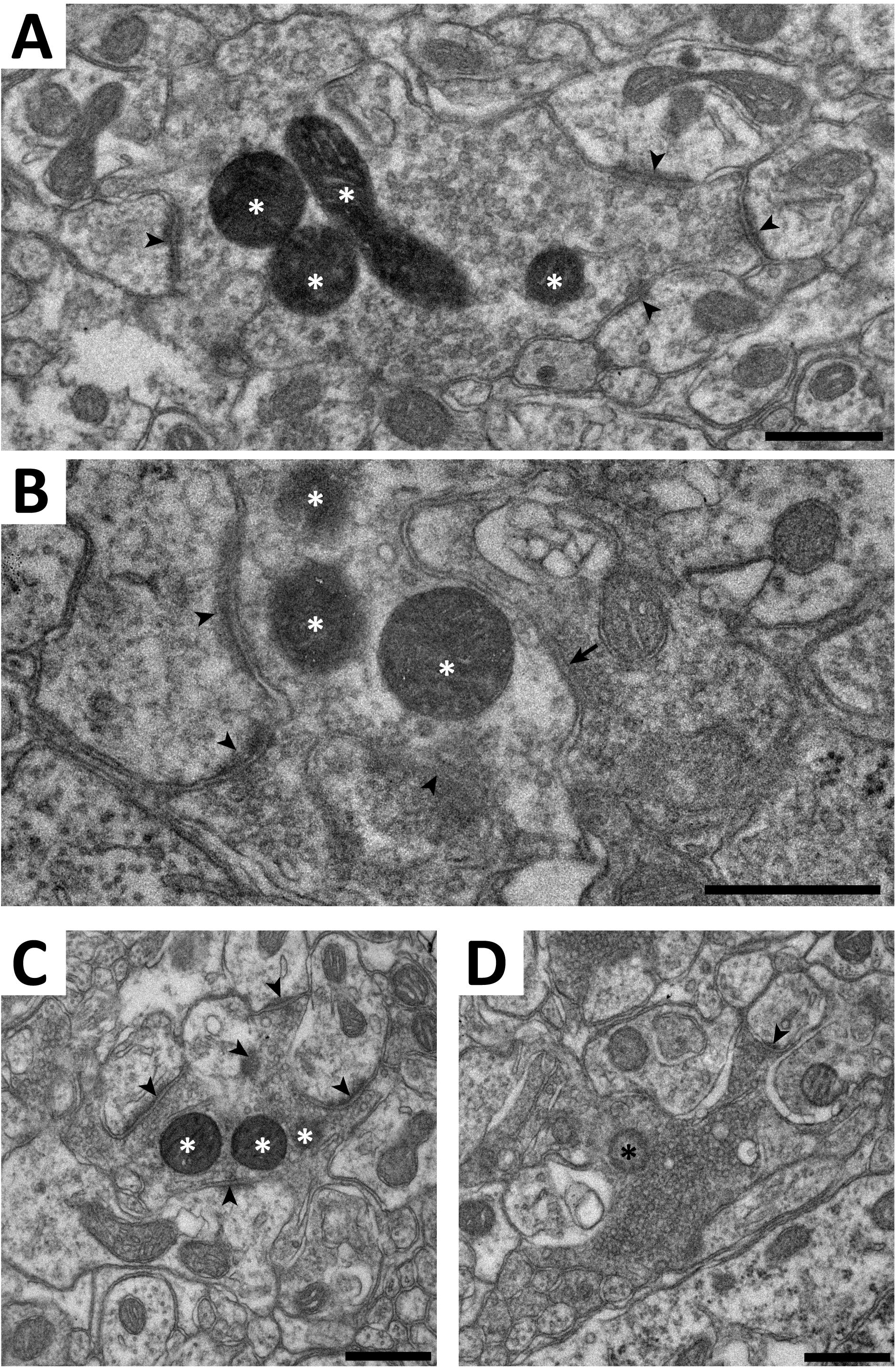
Electron microscopy of C-LTMR terminals in spinal cord of *Th*^CreERT2^;Na_V_1.8^FlpO^;APEX2 mice. **A-C**, examples of terminals forming synaptic type II glomeruli in lamina II. White asterisks indicate mitochondria with APEX2 reaction product. Arrowheads indicate asymmetric synapses established by the central APEX2^+^ terminal onto postsynaptic dendrites. Arrow in B indicates a symmetric presumed inhibitory synapse formed by a presynaptic axon onto the APEX2^+^ terminal. **D**, a type I glomerulus in mid-dorsal lamina II. The central terminal contain a mitochondrion (black asterisk) lacking APEX2 reaction product. Note the tightly packed synaptic vesicles of variable size, the electron dense axoplasm, and the scalloped shape, compared to the terminals in A-C. In particular, while the terminal in C has morphological characteristics resembling those in the terminal in D, they are much more pronounced in the latter. Scale bars, 500 nm.

## Discussion

In the present study we have generated and characterized a new knock-in mouse that expresses the recombinase FlpO from the locus of the gene *Scn10a*, that encodes the voltage-gated Na^+^ channel Na_V_1.8. We show that this mouse with very high efficiency and selectivity recapitulates the expression of Na_V_1.8 in primary afferent neurons, and can be used for intersectional targeting of primary afferent fiber populations, as exemplified here by the highly selective targeting of C-LTMRs by combining the new Na_V_1.8^FlpO^ mouse, presented in this study, with a *Th*^CreERT2^ mouse and double recombinase dependent reporter tools.

Na_V_1.8 was first identified in neurons of DRGs (Akopian et al. 1996) and found to be broadly expressed in nociceptors (Djouhri et al. 2003). A knock-in/knock-out Na_V_1.8^Cre^ mouse line was therefore developed (Stirling et al. 2005) which has since been widely used for nociceptor-specific targeting (e.g., Abrahamsen et al. 2008; Daou et al. 2013; Nassar et al. 2004; Lagerström et al. 2011). However, as Na_V_1.8 is broadly expressed in essentially all nociceptor populations identified, as well as in C-LTMRs (Kupari and Ernfors 2023; Usoskin et al. 2015), unmasking the roles of these subpopulations of sensory neurons is not possible using the Na_V_1.8^Cre^ mouse alone. Indeed, genetic markers are generally not selective for a single cell population but also expressed by other cell types, either in the PNS, CNS or non-neural tissue. Using Na_V_1.8 as one limb of an intersectional targeting approach is useful since it essentially restricts expression to primary nociceptors and C-LTMRs; another gene less specific to the PNS can then be used as a second driver of recombinase expression.

It has been reported that the previously available, widely used Na_V_1.8^Cre^ mouse line exhibits Cre-dependent recombination not only in nociceptors and C-LTMRs, but also in some A fiber LTMRs, including in nerve endings within Meissner corpuscles (Shields et al. 2012). Here we did not observe any recombined fibers innervating Meissner corpuscles, suggesting that our Na_V_1.8^FlpO^ mouse does not target any of the two types of rapidly adapting Aβ LTMRs that innervate Meissner corpuscles (Neubarth et al. 2020), in line with the expression of Na_V_1.8 mostly restricted to Piezo2-poor fibers as observed in transcriptomic studies (Sharma et al. 2020; Usoskin et al. 2015). Still, we did observe a similarly high proportion of recombined neurons that express the A fiber marker NFH as was found with the Na_V_1.8^Cre^ mouse line (∼40 %), suggesting that a substantial fraction of fibers exhibiting recombination were myelinated. Some reporter expressing innervation was also found in the dorsal column nuclei as well as in the principal nucleus of the trigeminal nerve, which would speak for FlpO expression in myelinated LTMRs. However, although dorsal column pathways are conventionally viewed as purely involved in low-threshold mechanoreception, they have also been suggested to be involved in visceral nociception, and this could be partly mediated by direct innervation by primary afferent branches (Willis et al. 1999). It can nevertheless, although FlpO-dependent recombination very closely followed the expression of the *Scn10a* (Na_V_1.8) gene, from the present observations not be excluded that a portion of the fibers showing such recombination were LTMRs.

We surveyed the distribution of recombined nerve fibers in a number of peripheral tissues. In nodose and jugular ganglia, many cells were tdTomato^+^, in line with previous reports using Na_V_1.8^Cre^ mice (Gautron et al. 2011; Stirling et al. 2005), as well as with transcriptomics data (Kupari et al. 2019). Accordingly, we consistently noted extensive innervation of vagal innervation targets, including the bladder wall, intestinal mucosa, and aortic wall. Numerous tdTomato^+^ fibers were present in the vaginal wall, including the mucosa and lamina propria, in accordance with the dense innervation by substance P^+^/CGRP^+^ fibers previously described (Barry et al. 2017). Similarly, the external and internal prepuces of the glans penis showed dense innervation of tdTomato^+^ fibers in alignment with recent observations of CGRP^+^ and TH^+^ fibers in these structures (Qi et al. 2024a).

A novel finding was the presence of recombined neurons in the Grüneberg ganglion, a little-studied sensory ganglion in the distal nose. This structure, which has been implicated in the sensation of alarm pheromones and cold (Brechbühl et al. 2008; Schmid et al. 2010), has previously been reported to not show Na_V_1.8 immunoreactivity (Schmid et al. 2010). Thus, the tdTomato expression found here could reflect either an only weak expression of Na_V_1.8, or ectopic recombinase expression.

Whereas much attention has been devoted to transcriptomic and neurochemical identification of cutaneous and visceral nociceptor populations, much less is known about nociceptor populations innervating skeletal muscle or other deep somatic tissue (Mense 2010). Here we found sparse but widespread innervation of recombined fibers in the gastrocnemius muscle, suggesting that the Na_V_1.8^FlpO^ mouse can be useful for characterizing of nociceptive muscle afferent fibers.

A somewhat surprising observation was the presence in the CNS of recombined cells and processes that did not originate from primary afferent sources. The presence of Na_V_1.8 protein or *Scn10a* mRNA in the brain has not been reported in the literature; however, the Allen Brain Atlas does show expression of *Scn10a* mRNA in several regions where we found recombined cells, including the granule cell layer of the olfactory bulb and layer 6 in parts of the frontal cortex. Further investigation is necessary to ascertain whether the reporter expression in our mice is attributed to actual Na_V_1.8 expression or to ectopic recombination.

We combined the Na_V_1.8^FlpO^ line with *Th*^CreERT2^ mice to test the utility of Na_V_1.8^FlpO^ mice in intersectional targeting, and demonstrated that this cross enables efficient and exquisitely selective targeting of C-LTMRs in DRGs without recombination in sympathetic neurons, Merkel cells or in other populations of DRG neurons. Notably, we found very little recombination in the trigeminal system, which is in agreement with the low expression of *Th* observed in C-LTMRs in trigeminal ganglia (Nguyen et al. 2017). A recent study generated a TAFA4^CreERT2^ mouse for targeting of C-LTMRs (Zhang et al. 2024), which selectively express this chemokine (Delfini et al. 2013). However, the TAFA4^CreERT2^ mice also showed recombination in subsets of DRG neurons with nociceptive or myelination markers, as well as in neurons in some regions of the CNS (Zhang et al. 2024). Thus, the intersectional *Th*^CreERT2^;Na_V_1.8^FlpO^ cross used here constitutes a more selective manner of targeting C-LTMRs.

While this manuscript was in preparation, the generation of another Na_V_1.8^FlpO^ mouse line was reported (Malapert et al. 2024). The distribution of recombined afferent fibers in spinal cord and peripheral organs in that mouse line is consistent with our observations. Malapert et al. used other reporter mice than those used here, precluding firm comparisons regarding efficiency and selectivity towards Na_V_1.8 expressing DRG neurons. However, Malapert et al. reported that about 85 % of *Scn10a*^+^ neurons expressed reporter (efficiency) and that 85 % of reporter-positive neurons expressed *Scn10a* (selectivity) in their mouse line, while we observed 97 % efficiency and 93 % selectivity in this study. Thus, it is possible that our newly generated mouse affords somewhat higher efficiency and selectivity than the previously reported strain.

In conclusion, here we report the generation of a Na_V_1.8^FlpO^ mouse line that enables a highly efficient and selective targeting of Na_V_1.8 expressing primary afferent neurons, as well as hitherto unrecognized small populations of such neurons in the brain. Furthermore, we show that this mouse line enables intersectional targeting of subpopulations of NaV1.8 expressing neurons such as C-LTMRs when combined with suitable Cre driver mice. We anticipate that this mouse line will be a highly useful and versatile tool for the continued exploration of murine sensory systems.

## Acknowledgements

This study was funded by Knut and Alice Wallenberg Foundation, project no. 2019.0047. We thank Dr. Maria Ntzouni and Dr. Vesa Loitto at the Microscopy Core Facility at the Medical Faculty, Linköping University for technical assistance.

