## Supplemental Text for "A NaV1.8^FlpO^ mouse enabling selective intersectional targeting of low threshold C fiber mechanoreceptors and nociceptors"

**Supplementary figure legends**

**Figure S1**. Co-localization of tdTomato and cell markers in DRGs from *Th*^CreERT2^;Na_V_1.8^FlpO^;Ai65 mice. Shown are example L3-5 DRG sections labeled for NeuN, CGRP, IB_4_ binding and NFH. Note lack of co-localization of tdTomato with the nociceptive markers CGRP and IB_4_ binding, as well as with the A fiber marker NFH. Scale bar is 50 µm, valid for all panels.

**Figure S2**. tdTomato^+^ fibers and cells in the brains of *Th*^CreERT2^;Na_V_1.8^FlpO^;Ai65 and *Th*^CreERT2^;Ai14 mice. Shown are parasagittal brain sections displaying complete lack of tdTomato fluorescence in the brain of *Th*^CreERT2^;Na_V_1.8^FlpO^;Ai65 mice except for the nucleus of the solitary tracy (Sol) and very sparse fibers in trigeminal areas (top panel), and the expected wide distribution of catecholaminergic structures in *Th*^CreERT2^;Ai14 mice (bottom panel). Scale bar is 1 mm, valid for both panels.
