## Supplementary figures and images for "A NaV1.8^FlpO^ mouse enabling selective intersectional targeting of low threshold C fiber mechanoreceptors and nociceptors"

### Supplemental Figure 1

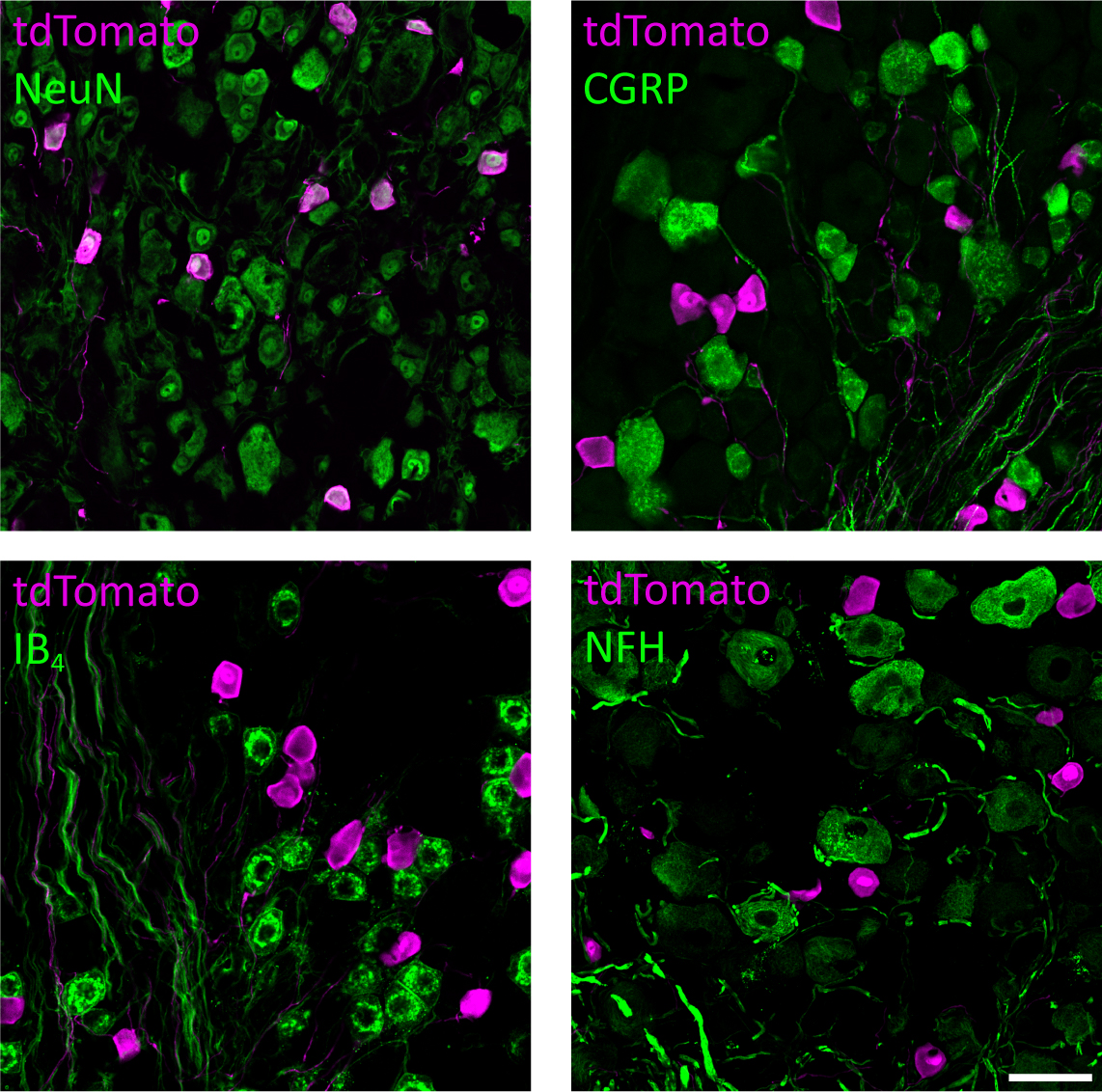

### Supplemental Figure 2

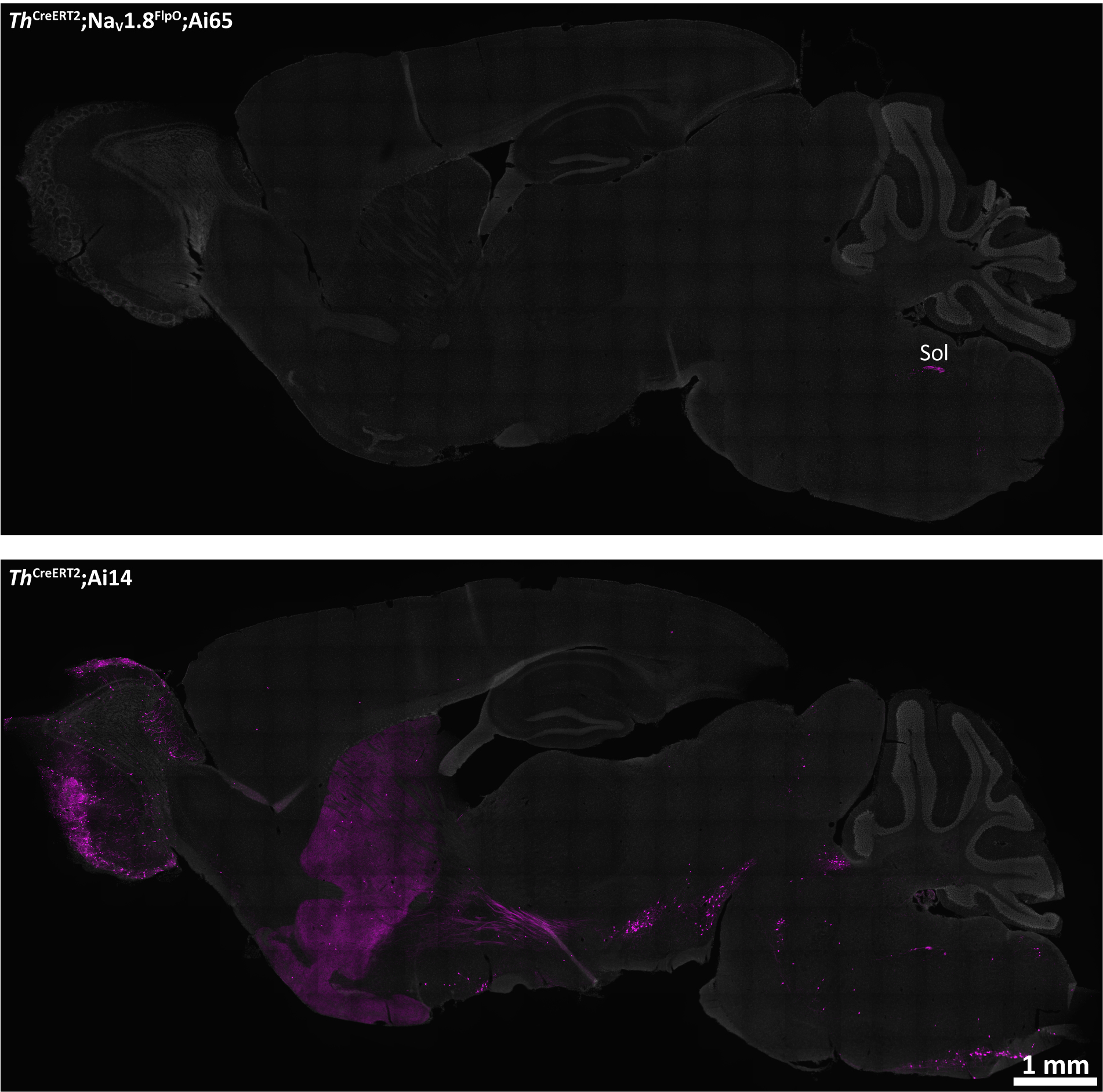
